# Molecular docking studies of potential inhibitors of acyl carrier protein and acetyl CoA Carboxylase in *Plasmodium falciparum*

**DOI:** 10.1101/2022.08.17.504218

**Authors:** Elliasu Y. Salifu, Issahaku A. Rashid, James Abugri, Festus Osei, Joseph Atia Ayariga

**Author notes:** Corresponding Author: James Abugri.

## Abstract

Malaria caused by *Plasmodium falciparum*, remains one of the most fatal parasitic diseases that has affected nearly a third of the world’s population. The major impediment to the treatment of malaria is the emergence of resistance of the *P. falciparum* parasite to current anti-malaria therapeutics such as Artemisinin (ART)-based combination therapy (ACT). This has resulted in countless efforts to develop novel therapeutics that will counter this resistance with the aim to control and eradicate the disease. The application of *in silico* modelling techniques has gained a lot of recognition in antimalarial research in recent times through the identification of biological components of the parasite for rational drug design. In this study we employed various *in silico* techniques such as the Virtual screening, molecular docking and molecular dynamic simulations to identify potential new inhibitors of biotin acetyl-coenzyme A (CoA) carboxylase and enoyl-acyl carrier reductase, two enzyme targets that play a crucial role in fatty acid synthesis in the Plasmodium parasite. Initially, 9 hit compounds were identified for each of the two enzymes from the ZincPharmer database. Subsequently, all hit compounds bind favourably to the active sites of the two enzymes as well as show excellent pharmacokinetic properties. Three (3) of the hits for the biotin acetyl-coenzyme A (CoA) carboxylase and six (6) of the enoyl-acyl carrier reductase showed good toxicity properties. The compounds were further evaluated based on the Molecular Dynamics (MD) simulation that confirmed the binding stability of the compounds to the targeted proteins. Overall, the lead compounds Zinc38980461, Zinc05378039, and Zinc15772056, were identified for acetyl-coenzyme A (CoA) carboxylase whiles zinc94085628, zinc93656835, zinc94080670, zinc1774609, zinc94821232 and Zinc94919772 were identified as lead compounds for enoyl-acyl carrier reductase. The identified compounds can be developed as a treatment option for the malaria disease although, experimental validation is suggested for further evaluation of the work.

## Background

One of the most human deadly parasitic disease is malaria which affects more than a third of the global population with annual report of roughly 200 million cases in the past decade [1]. In this regard, malaria has affected the livelihoods of large populations world-wide more than any other infectious disease. Another available data suggest an approximately half a billion individuals are infected with *Plasmodium spp*. globally, with annual mortality ranging from 1.5 to 2.7 million, with children seriously affected [2]. Recently, an estimated 241 million malaria cases have been reported world-wide in 2020, wherein, roughly 627, 000, up to 69 000 have been estimated to have died from the disease within previous year under review[1]. In view of the aforementioned alarming situation of the disease, there is urgent need to discover potent therapeutic targets for the development of new antimalarial drugs. This is because resistance to mainstream antimalarial drugs of *Plasmodium falciparum* impede the fight against malaria, which hamper the control and eradication strategies currently in place. Thus, the goal of eradicating the spread of malaria relies on finding therapeutic strategies against *P. Falciparum*.

The currently available antimalarial therapeutics as recommended by the World Health Organization include the Artemisinin (ART)-based combination therapy (ACT), antifolates, antimicrobials, Quinolines, etc[3], [4]. ACT is a combination of two or more drugs that work against the malaria parasite in different ways and used as a first line treatment. Also, a combination therapy of artesunate (AS) and amodiaquine (AQ) is another antimalarial therapy that has been in use for several years[5]. Despite the high potency and rapid action of these therapies in halting the spread of malaria, *P. falciparum* continues to find ways to resist being completely eradicated by acquiring resistance to the various available treatments[6]. Also, there is a reported high rate of recrudescence specifically associated with ART monotherapy or ACT[7]. The possible causes of recrudescence to these therapies is attributed to ART-induced ring-stage dormancy and recovery; although, little is known about the characteristics of dormant parasites. As such efforts are directed at finding new techniques to tackle the dormant state of the parasite to overcome this resistance[8].

To date, most research works are geared towards understanding the biology of the parasite including studying its genome with the aim of identifying crucial drug targets to develop novel therapeutics[9], [10]. In spite of the numerous efforts made, only a few targets have been confirmed through *in vivo* investigations and thus provide reliable leads for malaria therapy. An experiment by Chen et al, reported that most metabolic pathways are downregulated in dihydroartemisinin (DHA)-induced dormant parasites. However, the fatty acid and pyruvate metabolic pathways remain the only active mechanisms in the dormant parasites[11]. Thus the fatty acid synthetic pathway has been explored to be a crucial mechanism in the malaria parasite, hence possess a great potential for anti-malaria drug targets[11]. The biotin acetyl-coenzyme A (CoA) carboxylase and enoyl-acyl carrier reductase are two enzyme targets that play a crucial role in fatty acid synthesis in the Plasmodium parasite as reported by Chen et al., [11]. Particularly, these targets have been understood to interrupt recovery of the malaria parasites from ART-induced dormancy and reduces the rate of recrudescence after ART treatment[11]. However, despite their potentials in inducing the recovery of *P. falciparum* parasites’ from dormancy, the available literature suggests inconsistent results. Also, the current known IC50 of inhibitors that causes recovery of dormant parasites is deemed too high for their use as medicines. Thus the surge for potential new inhibitors is on the rise.

The use of molecular modelling methods is widely employed in antimalarial research through the identification of biological components of the parasite that can be targeted to develop novel therapeutics[12]. The discovery of these crucial enzyme targets (biotin acetyl-coenzyme A (CoA) carboxylase and enoyl-acyl carrier reductase) in the fatty acid synthetic pathway therefore provides a means for the identification of potential new inhibitors with improved efficacy than existing inhibitors of these targets using in silico modelling techniques [13]. Herein, we employed in-silico techniques including the in-house Per Residue Energy Decomposition (PRED)-based pharmacophore modelling, molecular docking, virtual screening and Molecular dynamic simulation to discover inhibitors of acyl-carrier protein and acyl-CoA carboxylase in *P. falciparum*. Additionally, the pharmacokinetic properties of all identified hit compounds were assessed which was followed by prediction of toxicity. We envision that findings from this study will form important basis for further experimental work to be carried out on these lead compounds to develop them into antimalarial agents.

## Computational Methodology

### Retrieval and preparation of protein and ligands

The 3D crystal structure of the biotin acetyl-coenzyme A (CoA) carboxylase and enoyl-acyl carrier reductase were retrieved from the RCSB Protein Data Bank[14] with respective PDB IDs: 1W96 [15] and 3F4B [16]. The two structures were experimentally solved through the X-ray diffraction method with a resolution value of 1.80°A and 2.49°A for 1W96 and 3F4B respectively. The 1w96 structure was retrieved in complex with soraphen A, a known inhibitor of acetyl-coenzyme A carboxylase, hence this was used as a reference compound in the study. Similarly, the structure of 3F4B was retrieved in complex with Triclosan which is a validated inhibitor of enoyl-acyl carrier reductase enzyme. As such this was also employed as reference compound in identifying potential inhibitors of enoyl-acyl carrier reductase. The retrieved structures and the reference inhibitors were prepared for a 20ns Molecular dynamic simulation to generate a Pharmacophore model at their stable states. Preparation for MD simulations was carried out on the graphical user interface of USCF chimera where all non-standard residues that are not relevant to the study were removed such as water, ions and other co-factors. In all, two systems were setup for a 20ns MD simulation comprising acetyl-coenzyme A in complex with Soraphen A (1W96) and enoyl-acyl carrier reductase in complex with Triclosan (3F4B).

### Pharmacophore Model Generation using Per Residue Energy Decomposition (PRED) Based Approach and Virtual Screening

The main goal in the Identification of small molecule antagonists of biotin acetyl-coenzyme A (CoA) carboxylase and enoyl-acyl carrier reductase is to model novel compounds that could also act in a similar or more potent inhibitory capacity towards the target protein as the identified reference inhibitors. Unlike other traditional pharmacophore modeling techniques, the validated in-house PRED method was used to outline the pharmacophoric features of the ligand-receptor in order to retrieve a more tailored potential hit/s [17], [18]. This pharmacophore model analyzes both the structural and chemical properties of proteins and ligands [17].To generate a PRED-based pharmacophore model, PRED decomposition was estimated via the MM/PBSA method for energy estimations after a 20ns MD simulations of the prepared complex systems. Pharmacophoric features based on the receptor-ligand interaction obtained from this short run MD simulation was selected. Residues Ile69, Lys73, Arg76, Ser77, Asn398, Val397, Gly396, Met393, Glu392 and Pro389 were found to be the highest energy contributing residues in the biotin acetyl-coenzyme A (CoA) carboxylase structure that interact with the ligand (Figure 1). Similarly, residues Ile333, Ala304, Ala305, Ile308, Val207, Gly204, Asn203, Ala202 and Tyr262 were identified as the highest energy contributing residues in the acyl carrier reductase complex (Figure 1). These identified moieties on each of the ligands that interacted with these residues were subsequently set as a query to generate a PRED-based pharmacophore models in ZINCPharma for the two targets of the fatty acid synthases pathways [19]. Subsequently, the zinc database was then screened for novel hits with similar features as the generated PRED-based pharmacophore models (Figure1)[20]. For hit screening, the filter was configured to query Zinc purchasable compounds with a molecular weight ≤500, with rotatable bonds set at ≤10. The “rule of five” proposed by Lipinski was also used as a cut-off [21]. Nine hit compounds were identified for each of the queried models on the ZincPharmer database[19]. The identified hits were downloaded in sdf file formats for further assessment.

**Figure 1:**
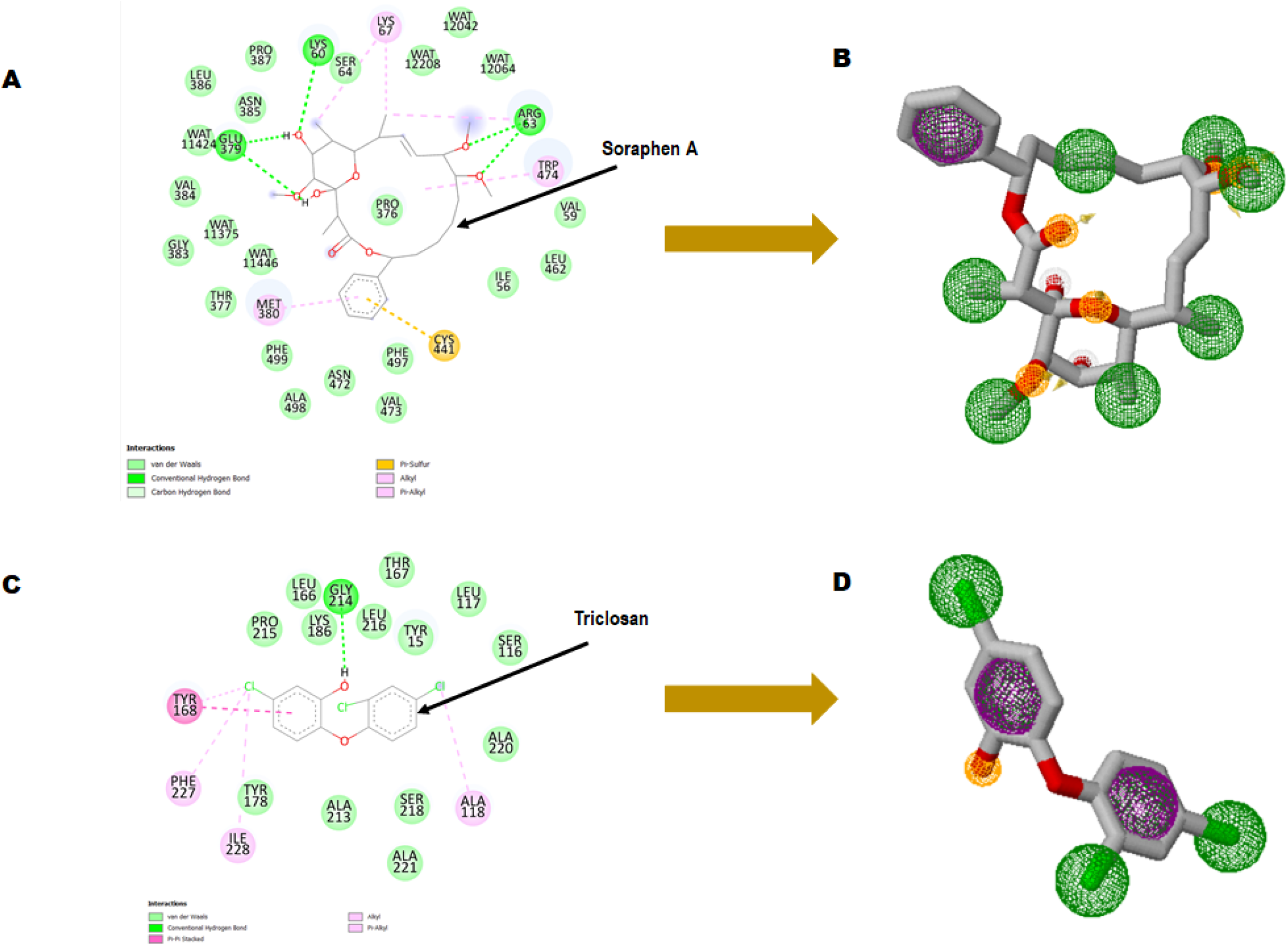
Ligand-residue interaction plot of (A) Soraphen A at the binding site of acetyl-coenzyme A (CoA) carboxylase enzyme and (C) Triclosan at the binding site of enoyl-acyl carrier reductase. (B) and (D) Generated pharmacophores showing aromatic (green) and hydrophobic moieties (Purple) and hydrogen donors (yellow).

### Molecular Docking of Hit Compounds

We conducted a molecular docking study on the identified hits to estimate the binding scores and also reveal the ligand-receptor best binding orientation between the hits and the acetyl-coenzyme A (CoA) carboxylase and enoyl-acyl carrier reductase. Molecular docking is also necessary to rank the hits in order of strongest binding affinity to the targets as this will screen out the most active hit compounds and yield a stable complex. Prior to molecular docking the hits compounds were taken through energy minimization using Avogadro 1.2.0 software[22]. The Avogadro software is incorporated with a UFF force field which optimizes the molecular geometries of the compounds and uses a steepest descent algorithm for structural minimization. The active site for each of the enzyme targets were determined using the grid positions of the native inhibitors (i.e., Soraphen A and Triclosan). The binding site based on the grid coordinates for acetyl-coenzyme A (CoA) carboxylase had the following coordinates; Centre (X=12.93, Y=26.86, Z=117.21) and Dimensions (X=25.99, Y=20.32, Z=21.16). The coordinates for the enoyl-acyl carrier reductase is as follow: Centre (X=-0.49, Y=-20.77, Z=-0.89) and Dimension (X=21.68, Y=20.32, Z=15.22). Subsequently, molecular docking was carried out using AutoDock vina incorporated in PyRx for all hit compounds[23], [24]. Output of docking was viewed on UCSF Chimera using the integrated ViewDock module after which the docking scores of the best pose for each complex were tabulated as illustrated in table 1 saved for further analysis.

**Table 1:** Binding Score of Hits via Molecular docking analysis.

| Acetyl Coenzyme A Carboxylase Hits |  |  |  | Enoyl-acyl Carrier Reuctase Hits |  |  |  |
| --- | --- | --- | --- | --- | --- | --- | --- |
| Index | Hit ID | Docking Score | RMSD Score(Å) | Index | Hit ID | Docking score | RMSD Score(Å) |
| 1 | ZINC05378039 | -7.3 | 1.51 | 1 | ZINC93658429 | -8.7 | 1.41 |
| 2 | ZINC38980461 | -8.1 | 1.73 | 2 | ZINC94085628 | -9.2 | 1.93 |
| 3 | ZINC38974815 | -8.3 | 1.81 | 3 | ZINC93656835 | -8.7 | 1.87 |
| 4 | ZINC38971181 | -8 | 1.99 | 4 | ZINC93098000 | -9.1 | 1.54 |
| 5 | ZINC38984088 | -8 | 1.33 | 5 | ZINC94919772 | -8.8 | 1.69 |
| 6 | ZINC38974798 | -7.4 | 1.89 | 6 | ZINC94080670 | -9.1 | 1.13 |
| 7 | ZINC38976811 | -8.2 | 1.51 | 7 | ZINC17074609 | -8.9 | 1.82 |
| 8 | ZINC15772056 | -9.8 | 1.23 | 8 | ZINC87263643 | -7.7 | 1.08 |
| 9 | ZINC38903582 | -7.9 | 1.67 | 9 | ZINC94821232 | -8.8 | 1.79 |
| 10 | Soraphen A | -12.1 | 1.77 | 10 | Triclosan | -7.5 | 1.88 |

### Assessment of pharmacokinetics properties Of Hits

In the computational drug design and development process, the early-stage assessment of pharmacokinetic parameters aids in the optimization of a molecular candidate to become an effective drug[25]. As such the resulting compounds obtained was evaluated and analyzed based on their physicochemical properties such as absorption, distribution, metabolism and excretion (ADME). Using the online platform SwissADME (http://www.swissadme.ch/index.php) [26], which helps to predict and analyse pharmacokinetic and pharmacodynamics properties of selected compounds. This was necessary to evaluate the prospects of the identified hits to be developed for human use. Furthermore, *In silico* ADME studies are expected to reduce the risk of late-stage attrition of drug development and to optimize screening and testing by looking at only the promising compounds[27]. ADME properties were predicted based on the Lipinski’s rule of five (LRo5). LRo5 is a general standard for estimating the biological activity, good oral bioavailability coupled with the tendency of a drug molecule to penetrate various aqueous and lipophilic (membrane) barriers [21], [28]. As all the identified hits were assessed for their drug-likeliness according the rule of five as shown in Table 2 and Table 3.

**Table 2:**
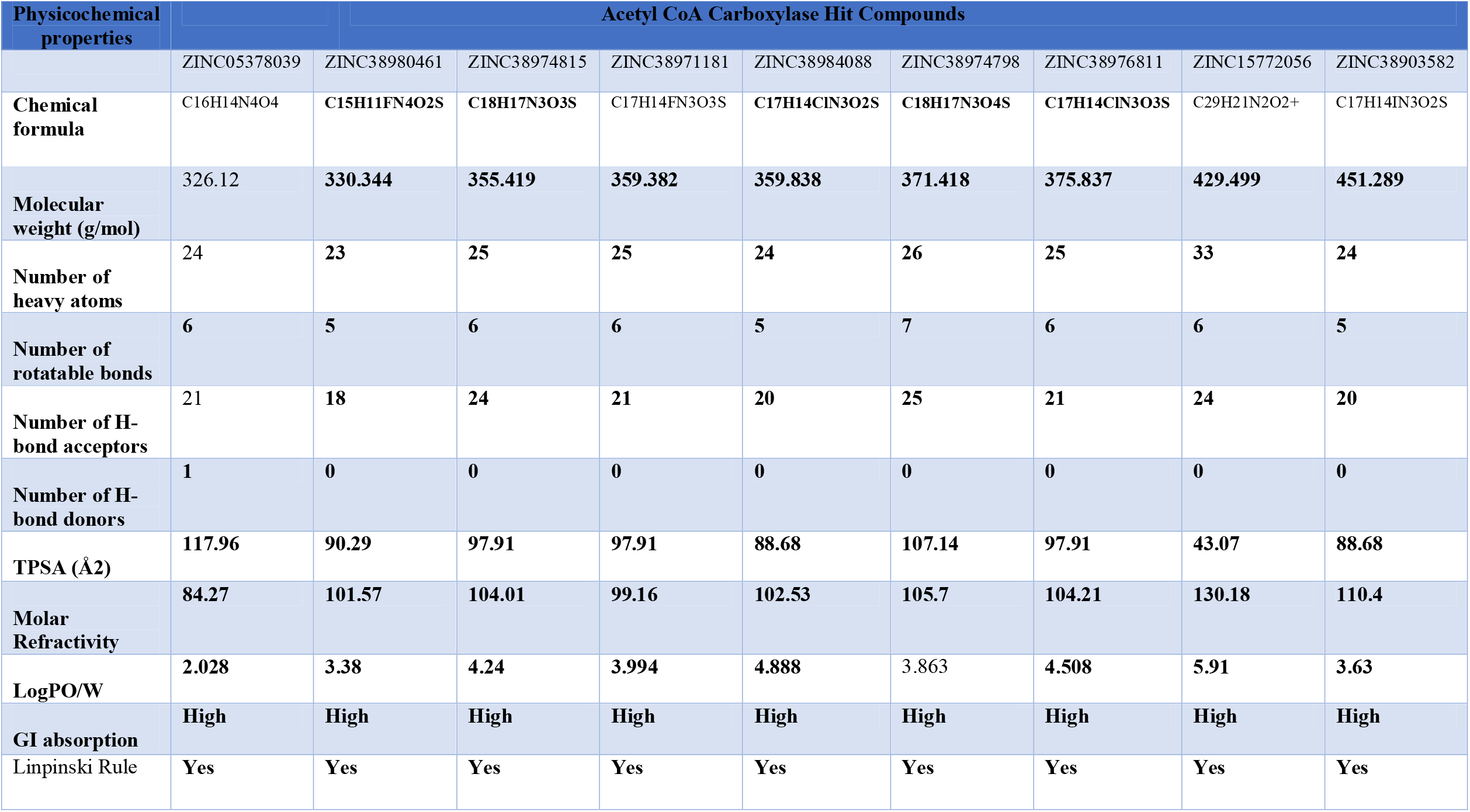
ADME properties of Acetyl CoA Carboxylase Hits

**Table 3:**
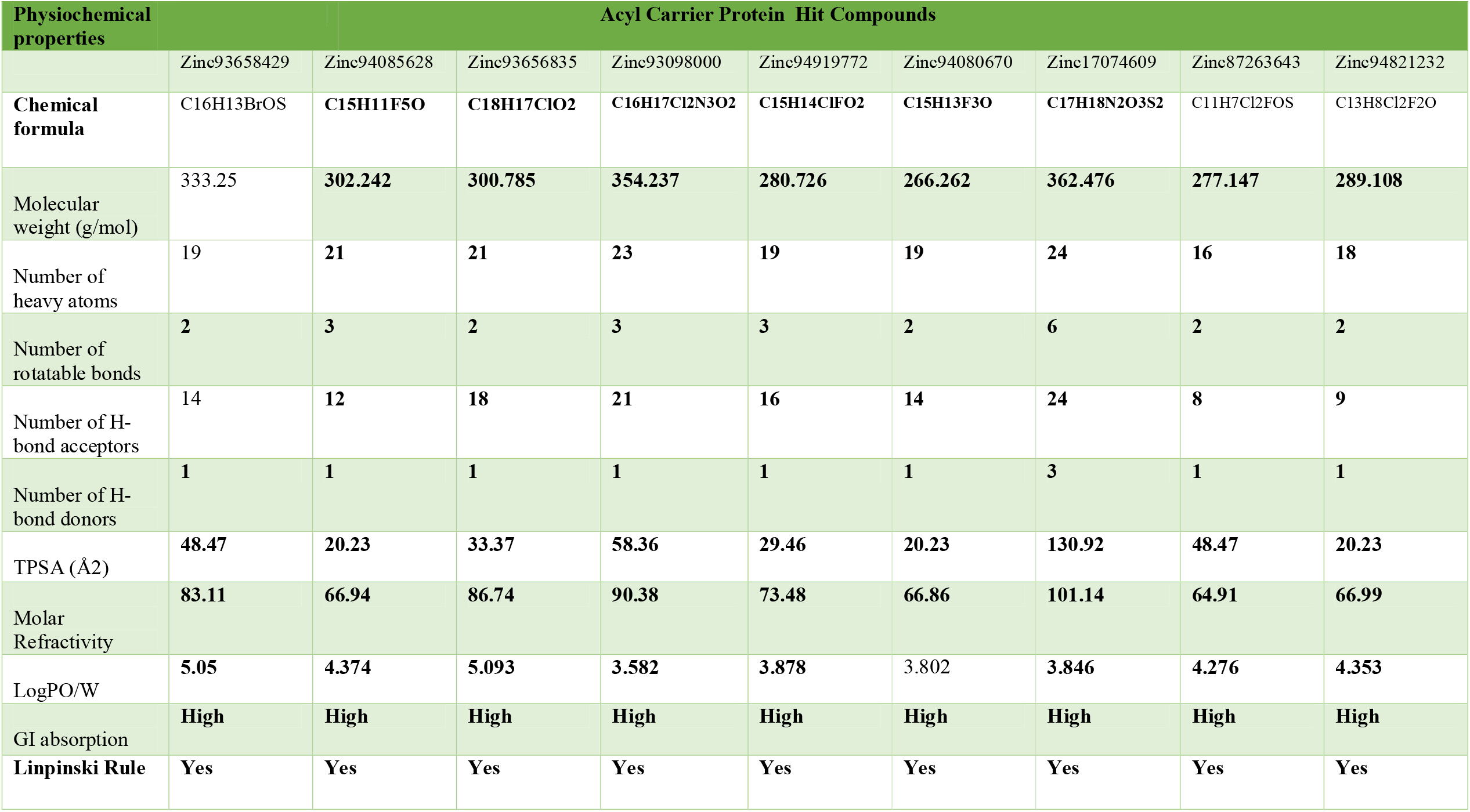
ADME properties of Acyl Carrier Protein Hits

### Prediction of toxicity

The prediction of chemical toxicity is a crucial step in the drug development process[29]. Not only are computational toxicity estimates faster in determining harmful levels in animals, but they can also assist to minimize the number of animal tests. Also, prioritizing compounds with a reduced risk of toxicity early in the drug development process should assist to reduce the high attrition rate in pharmaceutical R&D [29]. Chemists can be alerted if their suggested compounds are more likely to cause toxicity utilizing expert knowledge-based toxicity prediction. Toxicity prediction for the hit compounds was carried through ProTox-II (https://tox-new.charite.de/protox_II/) server [30]. Toxicity endpoints such as mutagenicity, carcinogenicity, and other characteristics can be quantified both quantitatively and qualitatively to determine a chemical compound’s toxicity using this server [31].

### Conformational stability Analysis of top hits via Molecular dynamic simulation

Molecular dynamic simulation was carried out for the top hits with good binding score, formed active interactions with targets, excellent pharmacokinetic properties and reduced toxicity effect. This is necessary to further analyse the impact of the selected hits on the stability and flexibility of the target enzymes. Receptor-Ligand structural assessment is crucial to the function of the receptor as any perturbation on the structural architecture of the protein/enzymes will ultimately alter its biological effect. All selected systems for MD simulation were prepared on UCSF Chimera comprising docked complexes of all nine (9) hits for each of the two targets, an unbound apo system for each of the targets and a reference system of soraphen A in complex with Acetyl CoA and triclosan in complex with Acyl carrier protein.

All molecular dynamics (MD) simulations were carried out using the graphic processing unit (GPU) version of the Particle Mesh Ewald Molecular Dynamics (PMEMD) of AMBER 18 software package [32].Atomic partial charges for the hit compounds were generated by the ANTECHAMBER module by using the Restrained Electrostatic Potential (RESP) and the General Amber Force Field (GAFF) protocol [33]. The receptors were parametized by the FF14SB [34] force field integrated in the Amber 18 suit. The Link Edit and parm (LEAP) module [35] of Amber 18 was then used to add hydrogens that are missing from the systems during preparation. Also, this module neutralizes the system by adding counter ions such as Na+ and Cl-after which the systems are solvated by suspending them in Transferable Intermolecular Potential with 3 Point (TIP3P) water box of size 8Å. A complexed coordinates and topology files of the receptor-ligand binding are generated for subsequent processing. The systems were minimized for 2000 energy steps. Initial minimization of 1000 steps with steep descent were performed for all the systems with a restrain potential and then followed by another 1000 steps minimization by conjugate gradient algorithm without restrain. The systems were then gradually heated from 0K to 300K with a 5kcal/mol. A harmonic restraint potential in NTP ensemble using Langevin thermostat of collision frequency of 1/ps. All the systems were then equilibrated at 300K for 500ps without restraint with a constant pressure at 1 bar using Berendsen barostat. SHAKE algorithm was used to restrain all hydrogen bonds [36]. MD production of 100ns was then performed without restrain on the systems with target coupling of 2 ps and constant pressure at 1 bar. Analysing the trajectories and coordinates generated from the MD run was carried through the CPPTRAJ and PTRAJ modules [37] incorporated in Amber 18. The Root Mean Square Deviation (RMSD), and Root Mean Square Fluctuation (RMSF) were calculated for all the systems. Discovery Studio version v19.10.18289 [38] and UCSF chimera were used to visualize the trajectories while Origin data version 6.0 tool [39] was used to plot all graphs.

### Binding Free Energy Analysis via MM/GBSA Method

The Molecular Mechanics/Generalized Born Surface Area (MM/GBSA) [40]–[43] method was employed in estimating the binding free energy for each of the inhibitor-bound systems. The binding free energy (Δ**G**_**bind**_) was calculated from the following equation:

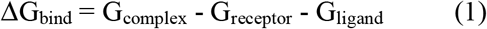

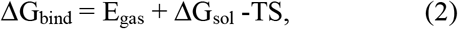

Where ΔG_bind_ is considered to be the summation of the gas phase and solvation energy terms less the entropy (TS) term

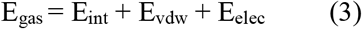

E_gas_ is the sum of the AMBER force field internal energy terms E_int_ (bond, angle and torsion), the covalent van der Waals (E_vdw_) and the non-bonded electrostatic energy component (E_elec_). The solvation energy is calculated from the following equation:

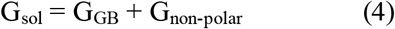

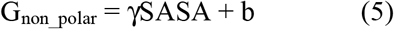

Where ΔG_bind_ is taken to be the sum of the gas phase and solvation energy terms less the entropy (TΔS) term., G_complex_ represents energy of the receptor ligand complex. Whiles G_receptor_ and G_ligand_ represents energies of receptor and ligand respectively. Egas denotes gas-phase energy; Eint signifies internal energy; and E_ele_ and E_vdw_ indicate the electrostatic and Van der Waals contributions, respectively. E_gas_ is the gas phase, elevated directly from the FF14SB force terms. G_sol_ denotes solvation free energy, can be decomposed into polar and nonpolar contribution states. The polar solvation contribution, G_GB_, is determined by solving the GB equation, whereas, G_SA_, the nonpolar solvation contribution is estimated from the solvent accessible surface area (SASA) determined using a water probe radius of 1.4 Å. T and S correspond to temperature and total solute entropy, respectively. γ Is a constant[44]. Per-residue decomposition analyses were also carried out to estimate individual energy contribution of residues of the substrate pocket towards the affinity and stabilization of each target.

## RESULTS

### Identification of Hit compounds

A pharmacophore structure defines how the key molecular properties of a ligand-receptor interaction are organized. As shown in figure 1, the top amino acids interacting with each of the targets are highlighted and subsequently, the pharmacophore model generated. The ZincPharmer database, a subsidiary of the zinc database was utilized to identify potential hit compounds based on the modelled pharmacophore. From our findings 9 hit compounds were identified for each of acetyl-coenzyme A (CoA) carboxylase and enoyl-acyl carrier reductase from the ZincPharmer query search (Figure 2). The hits were subjected to a molecular docking to assess their binding affinity to the target proteins.

**Figure 2:**
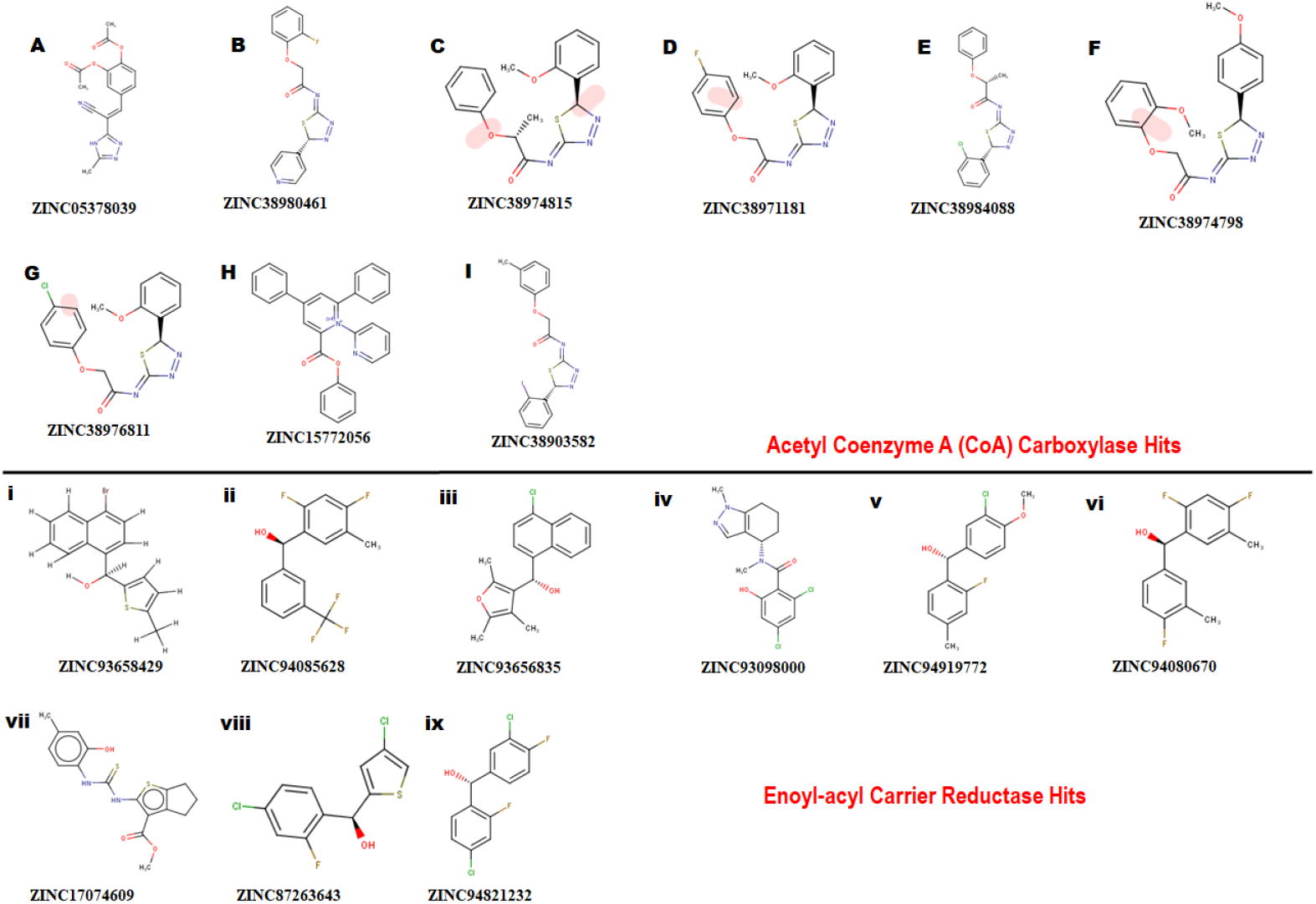
Showing 2D structures of identified hits. (A)-(I) comprising hits for Acetyl Coenzyme A (CoA) carboxylase and (i)-(ix) comprising hits for Enoyl-acyl carrier reductase.

### Analysis of binding score of Hits using Molecular Docking

Molecular docking was utilized extensively to predict the binding affinity and orientation of all screened hit compounds from the Zincpharmer when they bind to their biological targets. Our findings illustrated in table 1 show the respective binding scores of the hit compounds. Also, included are the assessment binding affinity scores for the reference compounds soraphen A and Triclosan used in the study. For most binding score analysis, the most negative value is usually indicative of a stronger binding and vice versa. In assessing the hits that target Acetyl Coenzyme A Carboxylase, the threshold of binding affinities was observed to be in the range -7.3kcal/mol and -9.8kcal/mol. Also, all hits bound to the active site with a low RMSD score below <2 which indicates stability. Which is indicative of favorable binding of all hits to the active site of Acetyl Coenzyme A Carboxylase. However, ZINC15772056 had the highest binding score of -9.8kcal/mol whiles ZINC05378039 had the lowest binding score among all the hits. The reference Soraphen compound showed to have the strongest binding score of -12.8kcal/mol as predicted from the molecular docking analysis. Similarly, all hit compounds of Enoyl-acyl carrier reductase bound favourably to the active site of their target which is evidenced by the binding score threshold in the range - 7.7kcal/mol and -9.2kcal/mol for all hit compounds. This finding shows that all screened hits bind favourably to Enoyl-acyl carrier reductase based on the molecular docking prediction. However, ZINC94085628 had the highest binding score of -9.2kcal/mol whiles ZINC87263643 had the lowest dock score of -7.7kcal/mol. Overall, all hit compounds showed to have bound strongly in contrast to the reference Triclocan compound with low RMSD score correlating to a stable complex.

### Assessment of ADME Properties of Hit compounds

Beyond the experimental models, determining drug-likeness reveals pharmacokinetic and pharmacodynamics aspects that unavoidably impact metabolism, distribution, absorption, and excretion in human systems. In predicting the drug-likeliness of a chemical compound, the Lipinski’s rule of five (LRo5) is considered. LRo5 is a universal benchmark for determining a drug’s biological activity, excellent oral bioavailability, and ability to permeate different aqueous and lipophilic (membrane) barriers[21], [45]. SwissADME [26] was used to predict physiochemical and pharmacokinetic properties of the screened compounds by following LRo5 thereby assessing their druggability. The descriptors, according to the LRo5 include molecular weight (MW) [≤ 500 Da], octanol-water partition coefficient [log P ≤ 5], Hydrogen bond donors (HBD) [≤ 5] and Hydrogen bond acceptors (HBA) [≤ 10]. As estimated from our findings in table 2 and table 3 all hits across both targets have molecular weights below 500Da threshold. A low molecular weight compound is mostly attributed to less toxicity and also an indication of a high tendency to be favoured for cellular uptake with little or no obstruction to its transport and distribution to target sites as opposed to compounds with larger MW. All other standards according to the LRo5 were matched by all identified hit compounds and hence passed the Linpinski’s assessment of druggability.

### Toxicity Assessment

The study of a compound’s toxicity is an important part of the drug development process. As such the toxicity characteristics of a potential drug candidate must be identified before it may enter clinical trials. Toxicities are usually explored through expensive, time-consuming and life-threatening animal studies, thus *in silico* toxicity estimates are a good option. Compounds are divided into toxicity classes based on the severity of their effects. Also, through a single or short-term exposure, toxicity can disrupt the synthesis of important enzymes in an organism, leading to the failure of a key organ. Sometimes the chemicals developed as medication candidates are toxin-like and damaging to other organs, causing organ toxicity, immunotoxicity, mutagenicity, and cytotoxicity in humans and animals. The toxicity of the compounds was predicted using ProTox-II, an online chemical toxicity prediction platform that incorporates molecular similarity, fragment propensities, and machine learning to predict toxicities [46]. The proTox-II server determines the toxic properties of compounds through the predicted median lethal dose (LD50) in mg/kg weight. Therefore, the toxicity of all identified hits were assessed in this study. Three (3) out of the nine (9) hits namely; ZINC05378039, ZINC38980461 and ZINC15772056 were identified to be in toxicity class IV for the acetyl coenzyme A carboxylase enzyme, indicative of non-toxicity and non-irritating. This finding highlights these three hits to be more favourable in terms of toxicity in contrast to the other hits.

Similarly, ZINC94085628, ZINC93656835, ZINC94080670, ZINC17074609, ZINC87263643, and ZINC94821232 for the enoyl – acyl carrier reductase enzyme, were in class IV, indicative of the excellent toxicity properties characterized by non-toxic and non-irritating properties. However, ZINC94919772 showed to be the most favourable in terms of toxicity and was identified to be in toxicity class V. Indicative of non-toxic, non-irritating, non-harmful when swallowed among others.

### Analysis of Conformational Dynamics of Protein-Ligand Stability via MD simulation

An assessment of the conformational dynamics was carried out for the two fatty acid targets in complex with their respective hits to unveil insights into structural alterations via MD simulation. The MD simulation was also carried out to validate the findings from molecular docking and to elucidate the energetic contributions of binding free energy. We employed post-MD analyses protocols, including; Root-mean-square deviation (RMSD), and Root-mean-square fluctuation (RMSF), analysis to provide insights on the structural impact of the hit compounds on acetyl-coenzyme A (CoA) carboxylase and enoyl-acyl carrier reductase. These post-MD protocols measure stability, and flexibility of the c-α atoms in the backbone of acetyl-coenzyme A (CoA) carboxylase and enoyl-acyl carrier reductase during the 100ns simulation.

### Structural stability

A 100-ns long MD trajectory was established to analyze the structural dynamics in the conformations of all systems. The overall protein convergence and stability of MD trajectories were determined based on RMSD, as shown in Figure 3. In the Acetyl-coenzyme A systems, convergence was attained early in the simulation after about 5ns. This was followed by steady atomic motions in all systems till the end of the simulation as shown in Figure 3A. None of the systems appeared to be unstable as shown by the plateau shape of atomic motions. Overall, the RMSD averages estimated for all the Acetyl CoA bound and unbound systems were 2.56Å, 2.09Å, 2.69Å, 2.42Å, 2.60Å, 1.88Å, 2.40Å, 1.77Å, 2.82Å, 1.85Å, and 2.43Å for apo, ZINC05378039, ZINC38980461, ZINC38974815, ZINC38971181, ZINC38984088, ZINC38974798, ZINC38976811, ZINC15772056, ZINC38903582, Soraphen A respectively.

**Figure 3:**
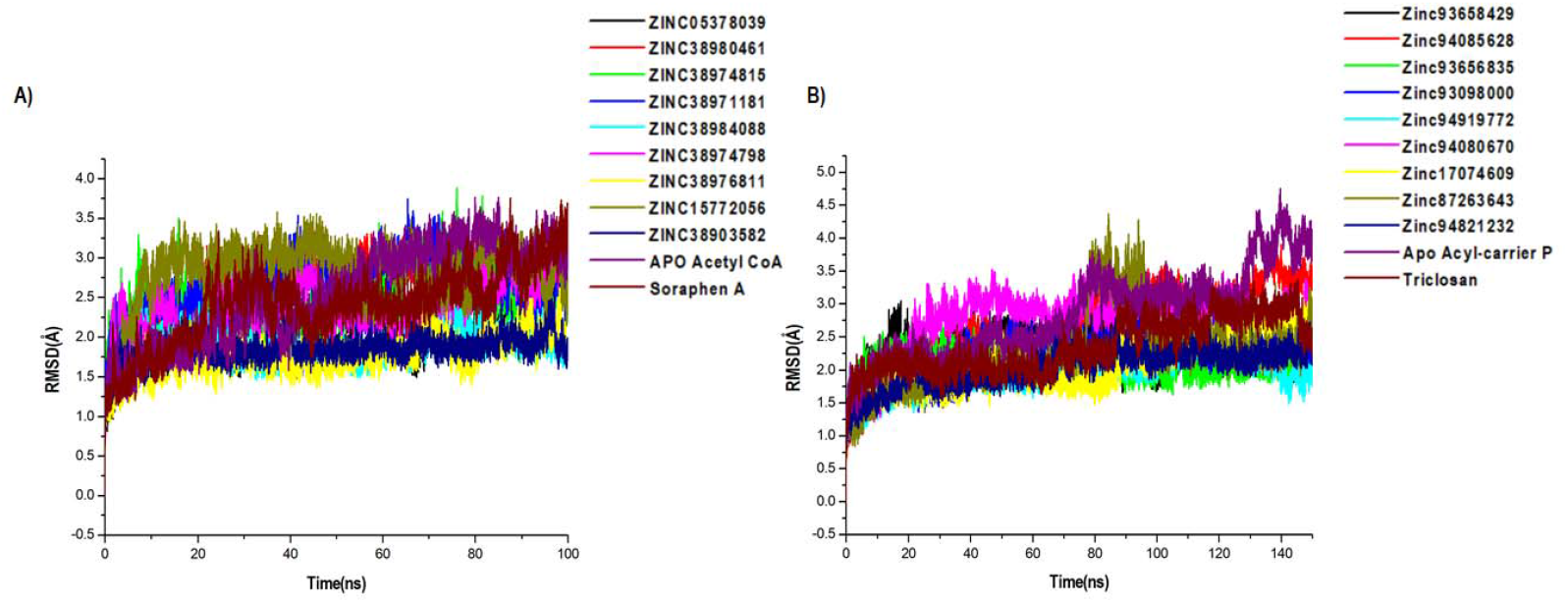
Comparative C-α RMSD plots showing the degree of stability and convergence of the hit compounds for (A) Acetyl coenzyme A carboxylase (B) Enoyl-acyl carrier reductase enzyme over the 100ns MD simulation time.

Similarly, all systems in the Acyl-carrier protein setup reached a convergence early in the simulation and maintained steady atomic motions of c-α atoms to the 100ns mark as shown in Figure 3B. All hit compounds including the unbound acyl carrier protein and the reference drug Triclosan appeared to be stable during the simulation. The estimated RMSD averages for all systems were 2.98Å, 2.28Å, 2.78Å, 2.10Å, 2.27Å, 1.95Å, 2.75Å, 2.23Å, 2.30Å, 2.08Å, and 2.45Å for Apo, ZINC93658429, ZINC94085628, ZINC93656835, ZINC93098000, ZINC94919772, ZINC94080670, ZINC17074609, ZINC87263643, ZINC94821232 and Triclosan respectively. These findings on structural stability of all systems highlight the reliability of our findings for further structural assessment.

### Structural flexibility

We employed RMSF analysis to determine the change in motion of each residue as a measure of the flexibility of certain regions of the acetyl-coenzyme A (CoA) carboxylase and enoyl-acyl carrier reductase structural architecture. A greater RMSF value typically indicates a more flexible structure, while a lower average RMSF value generally indicates a less flexible or rigid conformation. The average RMSF values estimated for all the Acetyl CoA bound and unbound systems were 1.23, 1.06Å, 1.20Å, 1.10Å, 1.02Å, 1.06Å, 1.05Å, 1.03Å, 0.98Å, 1.04Å and 1.37Å for the apo, ZINC05378039, ZINC38980461, ZINC38974815, ZINC38971181, ZINC38984088, ZINC38974798, ZINC38976811, ZINC15772056, ZINC38903582, Soraphen A respectively (Figure 4A). The average RMSF estimated for the Acyl-carrier protein systems were 1.50Å, 1.18Å, 1.42Å, 1.03Å, 1.01Å, 1.06Å, 1.21Å, 1.05Å, 1.10Å, 1.18Å and 1.13Å for Apo, ZINC93658429, ZINC94085628, ZINC93656835, ZINC93098000, ZINC94919772, ZINC94080670, ZINC17074609, ZINC87263643, ZINC94821232 and Triclosan respectively (Figure 4B). The effect of the hits on both fatty acid targets was evident as shown from the difference in structural fluctuations in the initial complexed structures and the structures after MD simulation as shown in Figure 4A and 4B.

**Figure 4:**
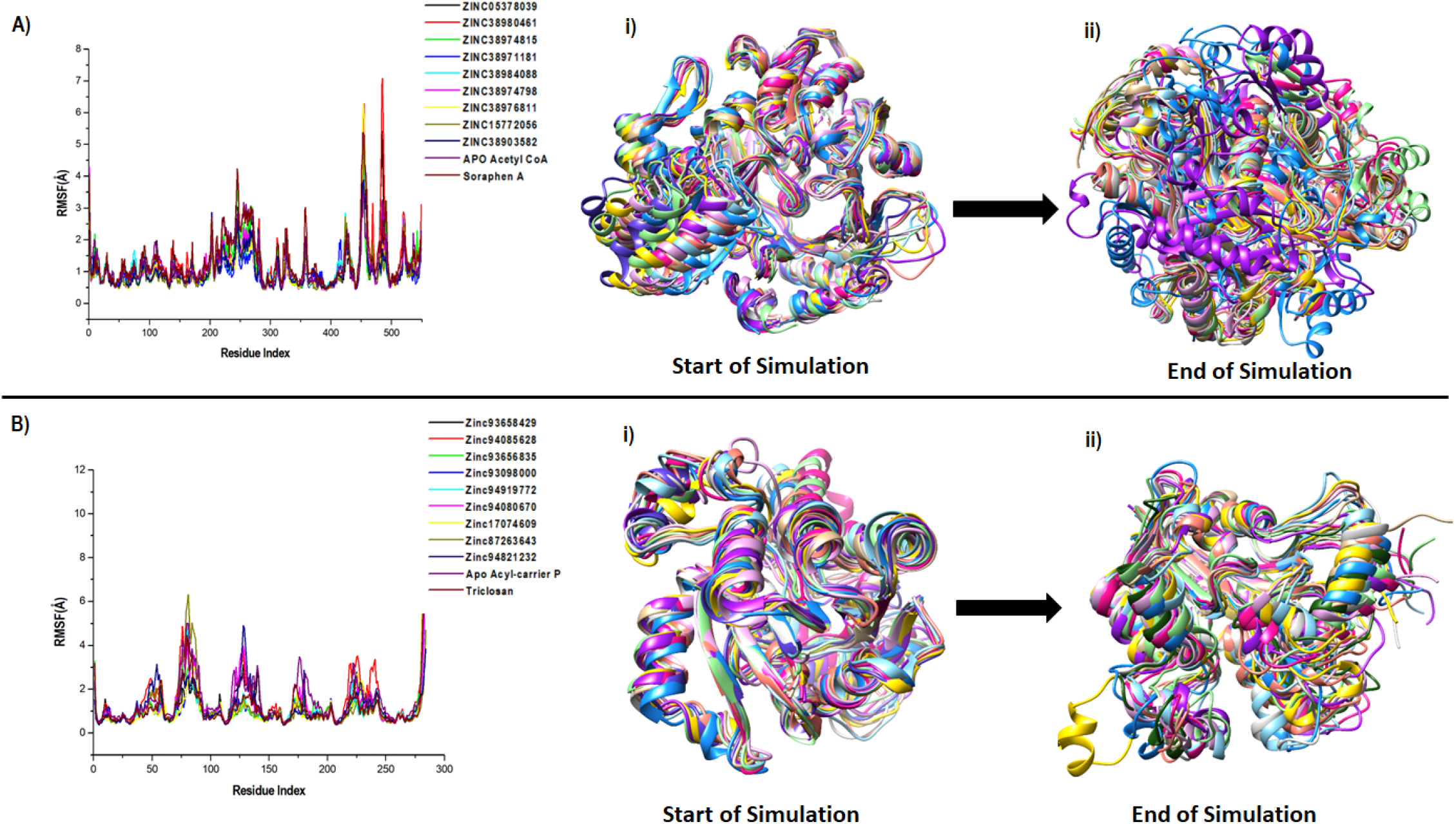
A plot of residual fluctuations in the unbound and bound **(A)** Acetyl co-enzyme A and **(B)** Acyl-carrier reductase enzyme. **i)** Highlights the initial superimposed structures of all systems, and **ii)** shows the various fluctuations that occurred at the end of the simulation.

### Binding Free Energy Assessment

The mechanics/generalized-born surface area (MM/GBSA) method was employed to estimate the binding free energetics of the bound complexes of the all hits including the two reference compounds Soraphen A and Triclosan. The molecular mechanics generalized Born surface area (MM/GBSA) is very popular method for binding energy prediction and is known to be more accurate than most scoring functions in molecular docking and are computationally less demanding than alchemical free energy methods [35]. The computed binding free energies of the acetyl-coenzyme A (CoA) carboxylase complexes ranged from - 15.78 to -36.36 kcal/mol whiles that of the enoyl-acyl carrier reductase complexes ranged from -32.89 to -41.42kcal/mol. Table 4 shows the energy terms that contribute to the binding free energy, the most favourable components being the Δ*E_ele_*, Δ*E_vdw_* and Δ*G*_gas_, while Δ*G*_sol_ was unfavourable. The energies presented by these compounds suggests the spontaneity, permeation and a measure of the reaction kinetics that characterize their complexing with the target proteins.

**Table 4:**
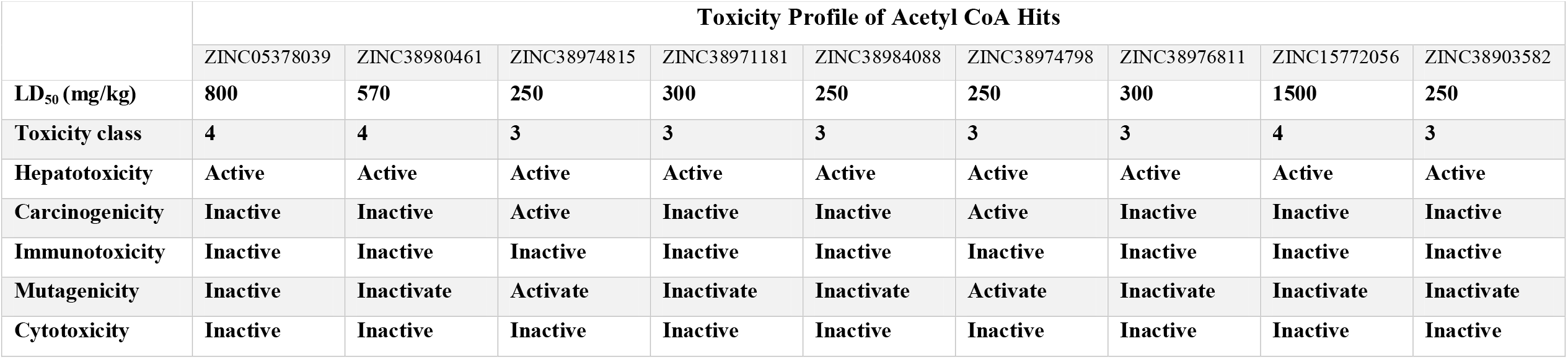
Toxicity assessment of Acetyl CoA Hits

## Discussion

Malaria still remains one of the mostly deadly parasitical disease in humans. A huge obstacle to the global efforts to control and eradicate malaria is the resistance of Plasmodium falciparum to conventional antimalarial therapies. As such several research studies have been geared towards identifying crucial therapeutics that will overcome this resistance. Advances in targeted therapy in antimalarial studies identified two crucial targets in the fatty acid synthetic pathway of the parasite. These new targets have been extensively investigated to interrupt recovery of parasites from ART-induced dormancy and to reduce the rate of recrudescence following ART treatment. The identification of these important targets have paved way for in-depth exploration into antimalarial therapy including the application *in silico* techniques in designing novel therapeutics that can pose an inhibitory effect to these targets.

In this study, we identified potential drug candidates against the acetyl-coenzyme A (CoA) carboxylase and enoyl-acyl carrier reductase of the fatty acid synthesis pathways using *in silico* techniques. Soraphen A and Triclosan, known inhibitors of these targets, were used as reference compounds in screening for potential drugs from the ZincPharmer database using the PRED Pharmacophore based virtual screening. This *in silico* technique allows the identification of moieties on the reference structures that form high affinity interactions with crucial amino acids residues on the target enzymes which form the basis for the generation of a pharmacophore. Subsequently, molecular docking was used to screen all hit compounds that were identified for each of the acetyl-coenzyme A (CoA) carboxylase and enoyl-acyl carrier reductase targets. All the 9 hit compounds of CoA showed favourable binding at the active site with good docking score and RMSD values < 2Å. Similarly, all the hit compounds in the acyl showed favourable binding as well with good binding score and lower RMSD score accounting for a stable binding. In most molecular docking studies an RMSD threshold <2Å is normally considered a good docking solution[47]. All our identified hits had RMSD less than the 2Å threshold.

Additionally, the pharmacokinetics was assessed by evaluating the ADME properties based on the Lipinski’s rule of five (RO5). The RO5 is used to demonstrate the drug-like properties of all selected compounds and serves as a justification for molecules that agrees with the rule. All the nine selected hits for both targets show good pharmacokinetic properties as well as passing the Linpinski’s test of “druggability”. Subsequently, the toxicity of all hit compounds was evaluated to unravel any harmful effects of the selected compounds on humans or animals. Three of the potential drug candidates for the Acetyl-coenzyme A target comprising Zinc05378039, Zinc38980461 and Zinc15772056 show the best toxicity properties characterized by being identified in the toxicity class iv indicative of no or less toxicity. They also show to be inactive towards Carcinogenicity, Immunotoxicity, Mutagenicity and Cytotoxicity effects. In the enoyl-acyl carrier reductase hit compounds, five (5) of the hits (zinc94085628, zinc93656835, zinc94080670, zinc1774609 and zinc94821232) were in class iv, indicating no or less toxic, however Zinc94919772 had the best overall toxicity properties which includes being in a toxicity class V correlating to no toxicity effects as well inactive towards Carcinogenicity, Immunotoxicity, Mutagenicity and Cytotoxicity.

Furthermore, molecular dynamic simulations was used to unveil the stability and flexibility of the selected ligands against the two fatty acid targets [48]. The cα atoms of the protein-ligand complexes were used to calculate the RMSD of the system that confirm low deviation of the system[48], [49]. Generally, the acceptable threshold for an average change in RMSD of the protein-ligand complex 1-3Å [47]. As such any RMSD value larger than the 1-3Å threshold indicates a vast conformational change in the protein structure hence unacceptable[47]. The RMSD averages for all the selected hit compounds in this study was within the 1-3 Å threshold which is indicative of good stability. The stability of the simulated systems highlights the reliability of our findings and further show the impact of the hit compounds on the targets. The fluctuation of the protein targets was also determined based on the RMSF value that also confirm averagely low fluctuations in all the inhibitor bound systems correlating to a less flexible protein structure.

In several drug design studies, the Molecular Mechanics Generalized Born Surface Area (MM-GBSA) approach has been used to accurately predict binding free energies[46], [50]. The estimated binding free energies of the Acetyl-coenzyme A in complex with all hit compounds ranged from -15.78 to -36.36 kcal/mol. Table 6 shows the energy terms that contributes to the total binding free energy, the findings show the major driving/favorable components to be Δ*E_elec_*, Δ*E_vdw_* and Δ*G*_gas_, while Δ*G*_sol_ was unfavourable. Similarly, in the enoyl-acyl carrier reductase complexes, the binding free energies were in the range -32.89 to -41.42kcal/mol. These energies were similarly driven by the Δ*E_elec_*, Δ*E_vdw_* and ΔG_gas_ energy components whiles the Δ*G*_sol_ term remain unfavourable. Zinc38980461 had the most favourable binding energy (-36.36kcal/mol) among the acetyl-coenzyme A hits whiles ZINC93098000 had the highest binding free energy (-41.42kcal/mol) in the enoyl-acyl reductase hit compounds. Overall, all hit compounds for both targets displayed favourable binding free energies towards the respective target which highlights their potential as inhibitors of these target proteins.

**Tabel 5:**
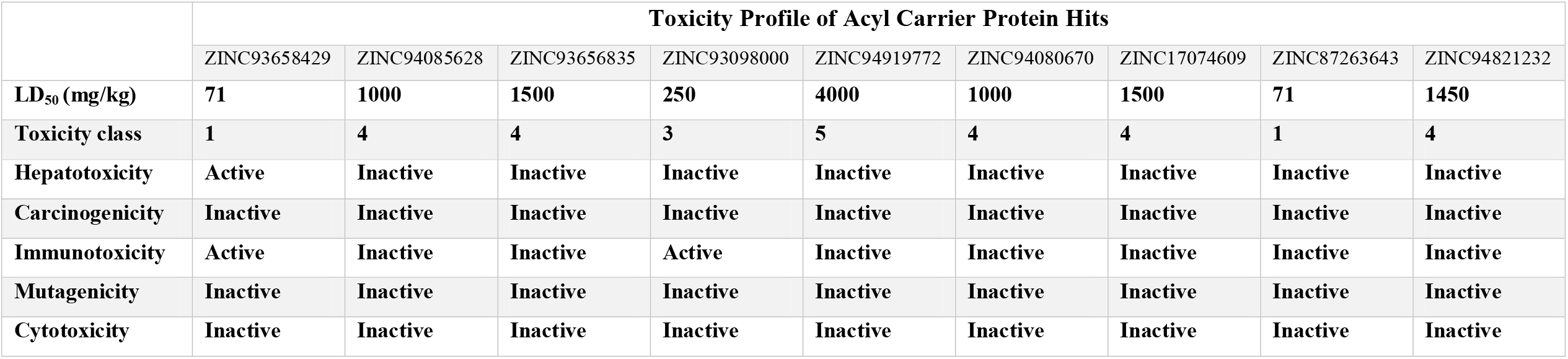
Toxicity assessment of Acyl Carrier Protein Hits

**Table 6.** MM/GBSA-based binding free energy profile of SdiA bonded terpenes and flavonoids.

| S/N | Acetyl CoA | Energy components (kcal/mol) |  |  |  |  | Acyl Carrier Protein | Energy components (kcal/mol) |  |  |  |  |
| --- | --- | --- | --- | --- | --- | --- | --- | --- | --- | --- | --- | --- |
| | | $\Delta E_{vdw}$ | $\Delta E_{ele}$ | $\Delta G_{gas}$ | $\Delta G_{sol}$ | $\Delta G_{bind}$ | | $\Delta E_{vdw}$ | $\Delta E_{ele}$ | $\Delta G_{gas}$ | $\Delta G_{sol}$ | $\Delta G_{bind}$ |
| 1 | ZINC05378039 | -25.1375 | -173.5097 | -198.6472 | 182.8632 | **-15.7840** | ZINC93658429 | -39.4309 | -16.4734 | -55.9043 | 17.9369 | **-37.9674** |
| 2 | ZINC38980461 | -37.4346 | -238.4188 | 244.1008 | 239.4950 | **-36.3584** | ZINC94085628 | -38.3190 | -8.6164 | -46.9336 | 12.1429 | **-34.7907** |
| 3 | ZINC38974815 | -33.2304 | -8.6508 | -41.8812 | 16.9848 | **-24.8963** | ZINC93656835 | -39.9459 | -6.9511 | -46.8970 | 11.6512 | **-35.2459** |
| 4 | ZINC38971181 | -36.0878 | -18.1971 | -54.2849 | 26.2825 | **-28.003** | ZINC93098000 | -41.4078 | -12.1751 | -29.2327 | 12.1902 | **-41.4229** |
| 5 | ZINC38984088 | -33.3977 | -20.2836 | -53.6813 | 23.0278 | **-30.6535** | ZINC94919772 | -40.2416 | -8.9398 | -49.1814 | 12.0991 | **-37.0822** |
| 6 | ZINC38974798 | -33.0974 | -13.5549 | -46.6524 | 23.6569 | **-22.6569** | ZINC94080670 | -37.7450 | -14.2537 | -51.9987 | 15.0559 | **-36.9428** |
| 7 | ZINC38976811 | -24.1298 | -14.3492 | -38.4791 | 21.3838 | **-17.0953** | ZINC17074609 | -42.0183 | -16.3969 | -58.4151 | 25.4525 | **-32.9626** |
| 8 | ZINC15772056 | -39.1807 | -382.7027 | -421.8833 | 392.4373 | **-29.4461** | ZINC87263643 | -35.6848 | -7.1093 | -42.7940 | 9.8949 | **-32.8991** |
| 9 | ZINC38903582 | -38.4375 | -7.3538 | -45.7913 | 14.3533 | **-31.4380** | ZINC94821232 | -36.8637 | -17.3503 | -54.2140 | 13.5729 | **-40.6411** |
| 10 | Soraphen A | -31.2378 | -11.5436 | -40.8371 | 113.4791 | **-18.2338** | Triclosan | -36.4314 | -3.8984 | -40.3298 | 6.9505 | **-33.3793** |

$\Delta E_{ele}$ = electrostatic energy; $\Delta E_{vdw}$ = van der Waals energy; $\Delta G_{bind}$ = total binding free energy; $\Delta G_{sol}$ = solvation free energy $\Delta G$ = gas phase free energy

Conclusively, based on our analysis of all hit compounds screened via the PRED-pharmacophore method for the acetyl-coenzyme A (CoA) carboxylase and enoyl-acyl carrier reductase of the fatty acid synthesis pathways, we identified Zinc38980461, Zinc05378039, and Zinc15772056 as the lead compounds that can be developed further into potential drug candidates of acetyl-coenzyme A, whiles zinc94085628, zinc93656835, zinc94080670, zinc1774609, zinc94821232 and Zinc94919772 were identified as the compounds with best properties against the enoyl-acyl reductase enzyme. The evaluations of the selected lead compounds were based on consistency in showing favourable results across different parameters including good docking score, excellent pharmacokinetic properties, no toxicity tendencies, ability to stabilize target enzyme with minimal fluctuations and a favourable binding free energy. These lead compounds can therefore be developed further into drug candidates for inhibiting acetyl-coenzyme A (CoA) carboxylase and enoyl-acyl carrier reductase of the fatty acid synthesis pathways in malaria therapy.

## Conclusion

The discovery of inhibitory molecules for a specific target protein is increasingly gaining attention in drug design due to the efficiency and speed that comes with the process. Using a computer-aided drug design methodology, we show in this study how novel acetyl-coenzyme A (CoA) carboxylase and enoyl-acyl carrier reductase inhibitors were quickly and effectively identified (CADD). Three lead compounds Zinc38980461, Zinc05378039, and Zinc15772056, were identified for acetyl-coenzyme A (CoA) carboxylase whiles zinc94085628, zinc93656835, zinc94080670, zinc1774609, zinc94821232 and Zinc94919772 were identified as lead compounds for enoyl-acyl carrier reductase, through the use of PRED based Pharmacophore method, virtual screening, molecular docking, ADMET, Toxicity assessment and MD simulation techniques in the CADD. These compounds may be able to inhibit the respective activities of acetyl-coenzyme A (CoA) carboxylase and enoyl-acyl carrier reductase and thereby interrupt the recovery of the falciparum parasite in host cell.

## Acknowledgements

The authors acknowledge, The Centre of High-Performance Computing (CHPC, www.chpc.ac.za), Cape Town, South Africa, for their resources in carrying out this work.

## Author Contributions

**Elliasu Y. Salifu**: Conceptualized the work, Contributed in the writing and editing of manuscript.

**Issahaku A. Rashid**: Assisted in setting up systems for MD simulations and proof reading of Manuscript.

**Festus Osei**: Contributed to writing of manuscript.

**Dr. James Abugri**: Supervised the work, assisted in writing and editing of work and final proof reading.

**Dr. Joseph Ayariga**: Proof read manuscript

